# Lipoate protein ligase B primarily recognizes the C_8_-phosphopantetheine arm of its donor substrate, and weakly binds the acyl carrier protein

**DOI:** 10.1101/2022.01.03.474776

**Authors:** Chetna Dhembla, Usha Yadav, Suman Kundu, Monica Sundd

**Affiliations:** National Institute of Immunology, Aruna Asaf Ali Marg, New Delhi 110 067, India; Department of Biochemistry, University of Delhi South Campus, Benito Juarez Road, New Delhi 110 021, India

**Author notes:** To whom correspondence should be addressed: Monica Sundd, National Institute of Immunology, Aruna Asaf Ali Marg, New Delhi-110 067, India. First two authors contributed equally to the work (Chetna Dhembla, Usha Yadav).

**Keywords:** Lipoate protein ligase B, Acyl carrier protein, Glycine cleavage system H protein, NMR, CoA, C_8_-CoA, lipoic acid synthesis, ACP

## Abstract

Lipoic acid is a sulfur containing cofactor, indispensable for the function of several metabolic enzymes. In microorganisms, lipoic acid can be salvaged from the surroundings by Lipoate protein ligase A (LplA), an ATP-dependent enzyme. Alternatively, it can be synthesized by the sequential action of Lipoate protein ligase B (LipB) and Lipoyl synthase (LipA), in a two-step reaction. LipB uptakes octanoyl-chain from C_8_-acyl carrier protein (C_8_-ACP), a byproduct of the type II fatty acid synthesis pathway, and transfers it to a conserved lysine of the lipoyl domain of a dehydrogenase. The molecular basis of substrate recognition by LipB is still not fully understood. Using *E. coli* LipB as a prototype, we show that the enzyme mainly recognizes the 4’-phosphopantetheine tethered acyl-chain of its donor substrate, and weakly binds the apo-acyl carrier protein. It can accept octanoate-from its own ACP, noncognate ACPs, as well as C_8_-CoA. Further, our NMR studies demonstrate the presence of an adenine and phosphate binding site in LipB, akin to LplA. A loop containing ^71^RGG^73^ sequence, analogous to the lipoate binding loop of LplA is also conserved. Collectively, our studies highlight commonalities between LipB and LplA in their mechanism of substrate recognition. This knowledge might be of significance in the treatment of mitochondrial fatty acid synthesis related disorders.

## INTRODUCTION

Lipoic acid is a hydrophobic cofactor, essential for the function of several metabolic enzyme complexes, *viz*. pyruvate dehydrogenase complex (PDHc) involved in pyruvate oxidation, 2-oxoglutarate dehydrogenase complex as a component of Krebs cycle (OGDHc), glycine cleavage system (GCS) in glycine degradation, branched-chain keto acid dehydrogenases (BCKDHc) in the metabolism of branched chain amino acids, and 2-oxoadipate dehydrogenase in lysine metabolism (1-5). These are multimeric enzymes with a conserved lipoyl domain, that serves as a substrate for lipoic acid modification. Lipoyl lysine acts as a “swinging arm”, helping to shuttle intermediates between the active site of multienzyme complexes(1,4,6-9). The flexibility of lipoic acid is crucial for substrate channeling and electron transfer during oxidation-reduction reactions (10).

Most of the knowledge of lipoic acid metabolism originates from studies conducted on *E. coli, L. monocytogenes, Bacillus subtilis* and *S. aureus*(11). In the presence of free lipoate or octanoate, lipoic acid is salvaged from the environment by lipoate-protein ligase A (LplA) (12-14). An activated lipoyl-5’-AMP intermediate is formed from ATP and lipoic acid on the surface of LplA. Thereafter, the ε-amino group of a lipoyl lysine attacks the noncovalently bound lipoyl-AMP, forming an amide linkage with the lipoyl moiety (3,8,9,14,15). Structurally, LplA comprises a large N-terminal catalytic domain (that binds lipoic acid), and a small C-terminal domain(16). In lipoic acid deficient environments, Lipoate protein ligase B (LipB), also known as Lipoyl-octanoyl transferase catalyzes the first biosynthetic step of lipoic acid synthesis(4). Studies on *M. tuberculosis* (PDB 1W66) and *T. thermophilus* (PDB 2QHS, 2QHT, 2QHU and 2QHV) LipB structures suggest a common structural scaffold, similar to the N-terminal catalytic domain of LplA. LipB functions as a cysteine/lysine dyad acyltransferase, cysteine 176 and Lys 142 of *M. tuberculosis* LipB (Cys169 and Lys 135 of *E. coli* LipB) functioning as acid/base catalysts(17-19). A covalent octanoyl-LipB thioester intermediate is formed by the transfer of octanoyl-chain from C_8_-ACP (acyl carrier protein) to Cys 176 of LipB(20). Subsequently, the thioester bond is attacked by the ε-amino group of the lipoyl lysine, resulting in C_8_-modification of the latter. In the next step, the octanoyl group tethered to the lipoyl domain is converted to lipoic acid by insertion of sulfur groups in the presence of LipA (lipoyl synthase).

In metazoans, three analogous enzymes participate in lipoic acid synthesis; Lipoyl (octanoyl) transferase 2 (LipT2) that transfers octanoyl chain from C_8_-ACP to the lipoyl subunit of glycine cleavage system, b) LIAS (lipoic acid synthase), a sulphur insertion enzyme that adds two sulphur atoms to the C_8_-chain to form lipoic acid, and c) LipT1, an amidotransferase, that relays octanoyl-chain/lipoic acid from the H subunit of Glycine cleavage system to the lipoyl carrier protein subunit of other dehydrogenases, *viz*. PDH, BCKDHc, αKGDH. Malfunction of any of these gene products causes lipoic acid synthesis defects in neonatal patients. Disorders range from defective mitochondrial energy metabolism, toxic levels of certain amino acids, neurological problems, and even death, depending on which lipoic acid synthesis gene malfunctions (11,21,22). Similar clinical phenotypes are also observed if the upstream mitochondrial FAS (mitochondrial fatty acid synthesis) does not function properly *e*.*g*. mitochondrial enoyl CoA reductase associated disorder (MECR) (23,24), malonyl-CoA synthetase (ACSF3), malonyl CoA-acyl carrier protein transacylase (MCAT) (25) *etc*. Defects in Glycine cleavage system gene also causes elevated glycine levels in the brain, leading to analogous neurological problems (26,27). Impaired dehydrogenase activity can also give rise to respiratory defects and muscle weakness due to the disruption of Krebs cycle (5,12,13).

Interestingly, all lipoic acid metabolism enzymes belong to the Pfam PF03099 family, share a similar scaffold despite low sequence conservation. They differ remarkably in their catalytic mechanism(20,28). Sufficient biochemical/ structural data is available for the free, as well as substrate bound forms of LplA from *E. coli, Thermoplasma acidophilum, Enterococcus faecalis, S. pneumoniae*, and *Mycoplasma hypopneumoniae*. However, the biosynthetic enzyme LipB is relatively less characterized, and its structure is available from two sources so far, *M. tuberculosis* and *T. thermophilus*. The mechanistic understanding of LipB-substrate interaction is also sparse, underscoring the need for in-depth studies. Two recent studies highlight the ability of LipB to allosterically distinguish between different acyl-ACP’s, *viz*. C_6_-, C_8_-, C_10_- and C_12_-ACP(29,30). Our studies extend their findings one step further, and show that the enzyme can discriminate between apo-, holo- and C_8_-ACP as well. LipB primarily recognizes the 4’-phosphopantetheine tethered octanoyl-chain of its donor substrate (C_8_-ACP), and weakly binds ACP per se. It can uptake octanoate from non-cognate ACP’s, as well as octanoyl-CoA. We further show that an adenine and phosphate binding site are conserved within its active site cavity. Interestingly, a mice LipT2 sample also successfully transferred C_8_-chain from C_8_-CoA to *E. coli* GcvH *in vitro*, suggesting a common mechanism of substrate recognition by all octanoyl transferases.

## RESULTS

Lipoate protein ligase B (LipB, UNP60720) catalyzes a bisubstrate reaction, using C_8_-ACP (Mol. wt 9.4kDa) as a donor substrate and lipoyl domain of dehydrogenases *e*.*g*. Glycine cleavage system H protein (GcvH), Pyruvate dehydrogenase complex (PDH), or 2-oxoglutarate dehydrogenase complex (OGDH) as octanoate acceptors(26). Limited structural/biochemical data on the LipB-ACP complex led us to carry out this in depth study. *M. tuberculosis* LipB 1.08Å resolution crystal structure (*Mt*LipB, PDB 1W66) has been used as a structural model to interpret our data, due to the absence of an *E. coli* LipB structure.

*M. tuberculosis* LipB comprises five central β-strands ↑β_1_,↑β_2_,↓β_3_,↑β_7_,↓β_6,_ enclosed by three long helices, *α*_2_ and *α*_3_ on one face, and *α*_1_ on the other side of β sheet, illustrated in Fig. 1A. Some of the catalytically important residues of *M. tuberculosis* LipB are shown as sticks, labeled cyan. The corresponding *E. coli* LipB residues are also labeled (black). A short stretch of beta strands comprising ↓β2’,↓β1’,↑β3’, called the ‘capping sheet’ is also present. Figure 1B shows the molecular surface of *Mt*LipB, colored based on coulombic charge, displaying the location of the active site cavity. Figure S1 compares the sequence of different octanoyl trasferases. *E. coli* LipB (UNP 60720) shares 32% sequence identity with *M. tuberculosis* LipB (UNP P9WK83, PDB 1W66), 28% with human mitochondrial LipT2 (UNP A6NK58), 29% with *Mus musculus* LipT2 and 26% with the *L. major* enzyme (UNP Q4A1B1), illustrated in Fig. S1.

**Fig. 1.**
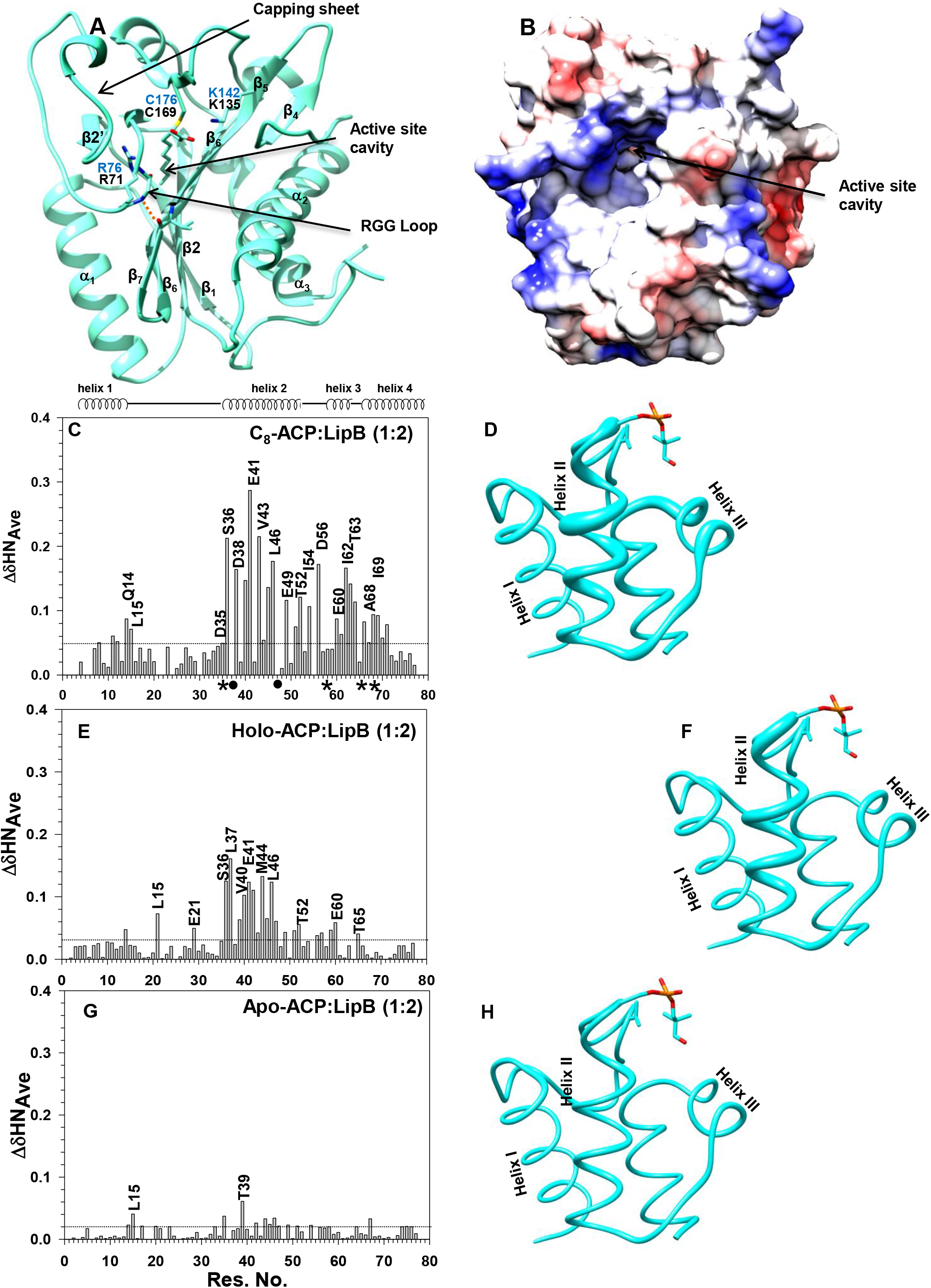
Chemical shift perturbations of *E. coli* ACP backbone upon LipB interaction. A) Ribbon representation of *M. tuberculosis* LipB (PDB 1W66) covalently bound to a decanoyl-chain. The positively charged residues in the active site cavity are shown as sticks. *M. tuberculosis* residue numbering is shown in blue, and the corresponding *E. coli* numbering in black. B) A surface representation of *M. tuberculosis* LipB, colored based on coulombic charge, displaying the opening of the active site cavity. Changes in the amide chemical shift of C) C_8_-ACP, E) holo-ACP, and G) apo-ACP, upon titration with unlabeled LipBK135A/C169A. ACP backbone has been represented as a worm (PDB 5VCB, chain D) for D) C_8_-ACP, F) holo-ACP, and H) apo-ACP upon LipBK135AC169A interaction. The magnitude of chemical shift change in each case is directly proportional to the thickness of the worm.

### Lipoate protein ligase B binds octanoyl-ACP with moderate affinity

To gain insights into the LipB: C_8_-ACP interaction, a catalytically inactive *E. coli* double mutant LipBK135A/C169A was prepared, and its interaction with ^15^N labeled *E. coli* C_8_-ACP was followed in a ^1^H^15^N TROSY-HSQC NMR experiment. ACP backbone was assigned based on the BMRB ID 27061, and standard 3D NMR experiments were acquired to assign the octanoyl-ACP spectra. Chemical shift changes upon LipB interaction have been reported as their weighted average, to give equal weights to proton and nitrogen chemical shifts, determined using equation 1.

*E. coli* ACP has a four helical bundle fold, comprising helix I (Thr 2-Gly 16), helix II (Asp 35-Asp 51), helix III (Asp 56-Thr 64) and helix IV (Thr 64-His 75). Upon titration with LipB, a number of backbone amides of C_8_-ACP present in helix II, III and IV, displayed significant perturbations (Fig. 1C). These include Gln 14, Leu 15, Asp 35, Ser 36, Asp 38, Val 40, Glu 41, Val 43-Leu 46, Glu 49, Thr 52, Ile 54, Asp 56, Glu 60-Thr 64, Glu 66, Ala 68-Tyr71. The peaks for Asp 35, Glu 57, Val 65 and Ala 68 were broadened beyond 1:1 ACP: LipBK135A/C169A ratio, shown by a (*) in the figure. Leu 37, and Glu 48 displayed chemical exchange line broadening right from the first titration point, shown as (•) in the figure. Perturbations have been mapped to the 1.1Å *E. coli* holo-ACP structure (PDB 5VCB, chain D) represented as a worm in Fig. 1D. The thickness of the worm is directly proportional to the magnitude of chemical shift change. Fig. S2A shows the multiple overlaid ^1^H^15^N TROSY-HSQC spectra for C_8_-ACP: LipBK135A/C169A titration. Each titration point is shown in a different color; free C_8_-ACP is colored red; C_8_-ACP:LipBK135A/C169A 1:0.25 molar ratio blue; 1:0.5 magenta; 1:0.75 cyan; 1:1 green, 1:1.5 orange and 1:2 purple. Same color coding has been used throughout, for all the titrations reported in this study.

The binding affinity of LipBK135A/C169A for C_8_-ACP was determined by Surface Plasmon Resonance (SPR) measurements, immobilizing LipB on the surface of a sensor chip and passing increasing concentrations of C_8_-ACP over the chip surface. Fig. S3A shows the SPR sensorgram, and S3B binding curve for C_8_-ACP: LipBK135A/C169A interaction. The two proteins interact in the micromolar range, with a dissociation constant (K_D_) of 5.5± 0.03 μM.

### LipB binds holo-ACP with comparatively lower affinity

Holo-ACP has a phosphopantetheine modification akin to C_8_-ACP, but lacks the octanoate-group. The contribution of the C_8_-chain in the interaction of LipB with its donor substrate was also investigated. ^15^N labeled holo-ACP (^15^N labeled 4’-phosphopantetheine arm) was titrated with unlabeled LipBK135A/C169A. Noticeable changes in the amide chemical shift were observed for several holo-ACP residues. As shown in Fig. 1E, Gln 14, Glu 21, Val 29, Ser 36, Leu 37, Thr 39, Val 40, Glu 41, Leu 42, Val 43, Met 44, Ala 45, Leu 46, Glu 47, Asp 51, Thr 52, Glu 53, Ile 54, Glu 57, Ala 59 and Glu 60 displayed a significant change. The aforementioned residues are present in helix II and III of ACP, also shown as a worm representation in Fig. 1F. Figure S2B displays superimposed multiple ^1^H^15^N TROSY-HSQC spectra, *i*.*e*. holo-ACP with increasing concentration of LipBK135A/C169A. Peaks that display significant change are shown by an arrow. Interestingly, one of the two NH resonances of 4’-Phosphopantetheine moiety (Phosphopantetheine ^15^N labeled) observed at a chemical shift of 8.1 ppm in the ^1^H^15^N TROSY-HSQC spectra, also displayed significant change in chemical shift during the titration (Fig. S2D).

Fig. S3C shows the SPR sensorgram and Fig. S3D binding curve for the interaction of LipBK135A/C169A with holo-ACP. A K_D_ of 31.9 ± 0.2 μM was obtained for the interaction.

### apo-ACP weakly interacts with LipB

In the cell, ACP is expressed as an apo-protein, and post-translationally modified to holo-ACP by the catalytic action of 4’-Phosphopantetheinyl transferase. The importance of the phosphopantetheine arm in the interaction of LipB with its substrate was also probed. ^1^H^15^N apo-ACP was titrated with unlabeled LipBK135A/C169A (Fig. 1G). Insignificant changes in amide chemical shift were observed (<0.06 ppm) upon addition of up to 2 molar equivalents of LipBK135A/C169A. In the multiple overlaid spectra, most of the apo-ACP amide resonances overlapped with the ACP peaks in presence of LipBK135A/C169A (1:2 molar ratio, Fig. S1C). A worm figure of ACP is also shown, displaying insignificant changes observed upon interaction with LipBK135A/C169A (Fig. 1H). Due to the weak nature of the complex, the interaction could not be probed by SPR.

### Non-cognate ACPs can serve as a donor substrate for LipB

Cross reactivity of LipB with non-cognate C_8_-ACP substrates was also investigated, as the enzyme displays weak affinity for its cognate ACP. The enzyme assay comprised *E. coli* LipB (5μM), an octanoate-donor (C_8_-ACP, 75μM), and an octanoate acceptor GcvH (25μM). Three separate assays were performed, using different octanoate-donors, a) *E. coli* C_8_-ACP (UNP 0A6A8), b) C_8_-*Lm*ACP (*L. major* ACP, UNP E9AD06), and c) C_8_-*Pf*ACP (*P. falciparum* ACP, UNP Q7KWJ1). The amount of product formed (C_8_-GcvH) in each case was qualitatively analyzed on a 12% SDS-PAGE gel, and quantitatively by C18-reversed phase chromatography. Figure 2A displays an SDS-PAGE gel for the product formed during the LipB assay. Lane 1 is pure apo-GcvH, Lane 2: C_8_-ACP, Lane 3: C_8_-*Lm*ACP, Lane 4: C_8_-*Pf*ACP (used as a control), and lanes 6-8 LipB assay carried out using C_8_-*Ec*ACP, C_8_-*Lm*ACP and C_8_-*Pf*ACP, respectively as donor substrates. Apo-to C_8_-GcvH conversion was discerned from the forward migration of the GcvH band on SDS-PAGE gel. The extent of GcvH conversion was also quantitatively analyzed by loading the samples on a C18 reversed phase HPLC column, and integrating the area under the elution peak, shown in Fig 2B-D. Approximately 90% GcvH conversion was observed using C_8_-ACP (*E. coli*) (Fig. 3B), 74% with C_8_-*Lm*ACP (*L. major* ACP) (Fig. 3C) and 85% in presence of C_8_-*Pf*ACP (*P. falciparum* ACP) as an octanoate-donor, illustrated in Fig. 2D.

**Fig. 2.**
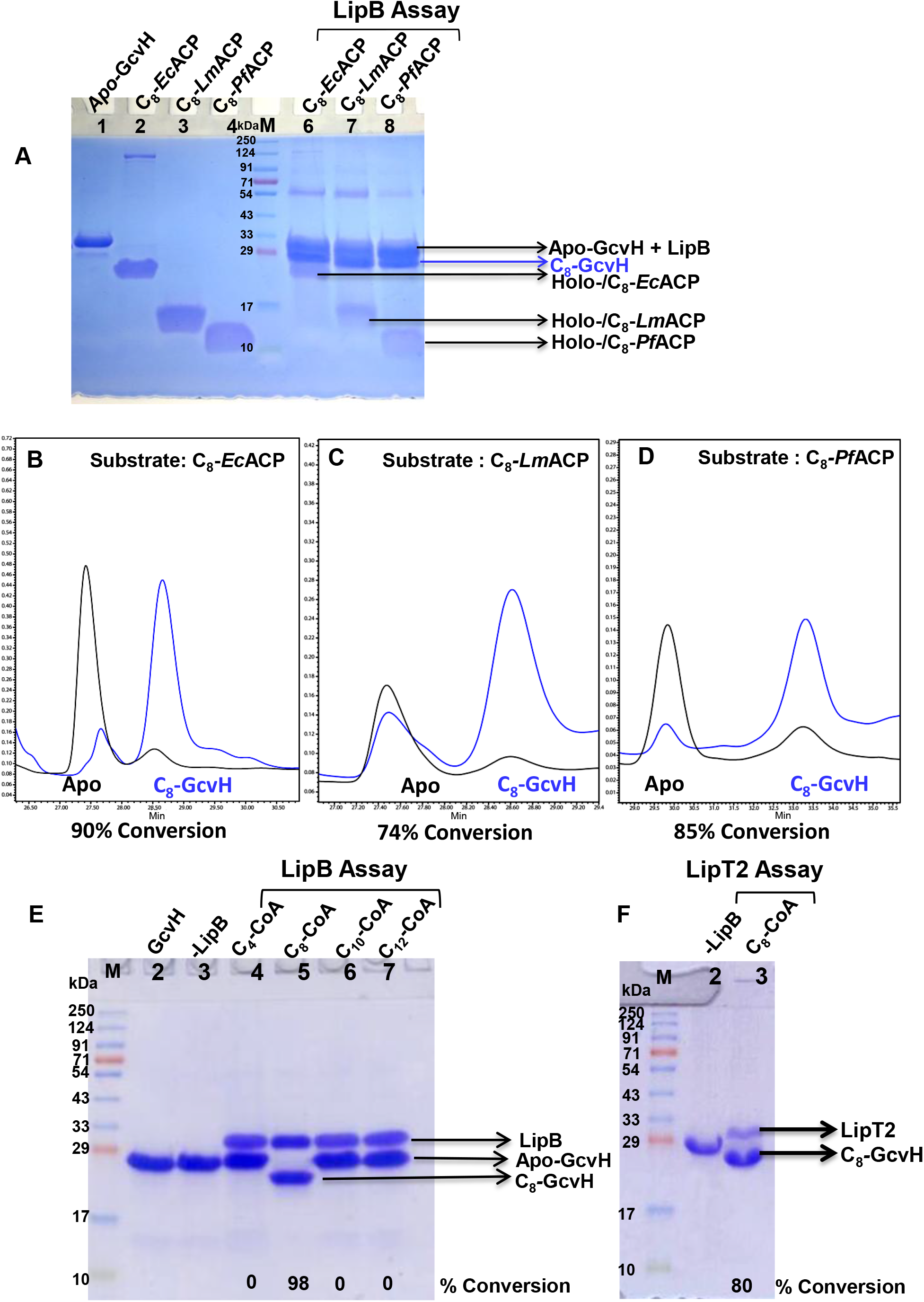
LipB can use non-cognate ACP as well as C_8_-CoA as octanoate donor. A) A 12% SDS-PAGE gel displaying, Lane 1: Apo-GcvH, Lanes 2-4: C_8_-ACP, C_8_-*Lm*ACP, and C_8_-*Pf*ACP loaded as control, Lane 5 Molecular weight marker, Lane 6-8: LipB assay using C_8_-ACP (*E. coli*), C_8_-*Lm*ACP (*L. major*), and C_8_-*Pf*ACP (*P. falciparum*), respectively as substrates. C18-reversed phase chromatogram for the LipB assay carried out using B) C_8_-ACP, C) C_8_-*Lm*ACP, and D) C_8_-*Pf*ACP, as octanoate donors. The chromatogram before the assay is shown in black, and after the assay in blue. %GcvH conversion after the assay was calculated and is shown at the bottom of each chromatogram. E) A 12% SDS-PAGE gel, displaying the conversion of GcvH, using different acyl-CoA as octanoate-donors. Lane 1: apo-GcvH, Lane 2: assay in the absence of LipB, Lane 3-6: assay in presence of C_4_-CoA, C_8_-CoA, C_10_-CoA and C_12_-CoA, respectively. GcvH band migrated faster in lane

**Fig. 3.**
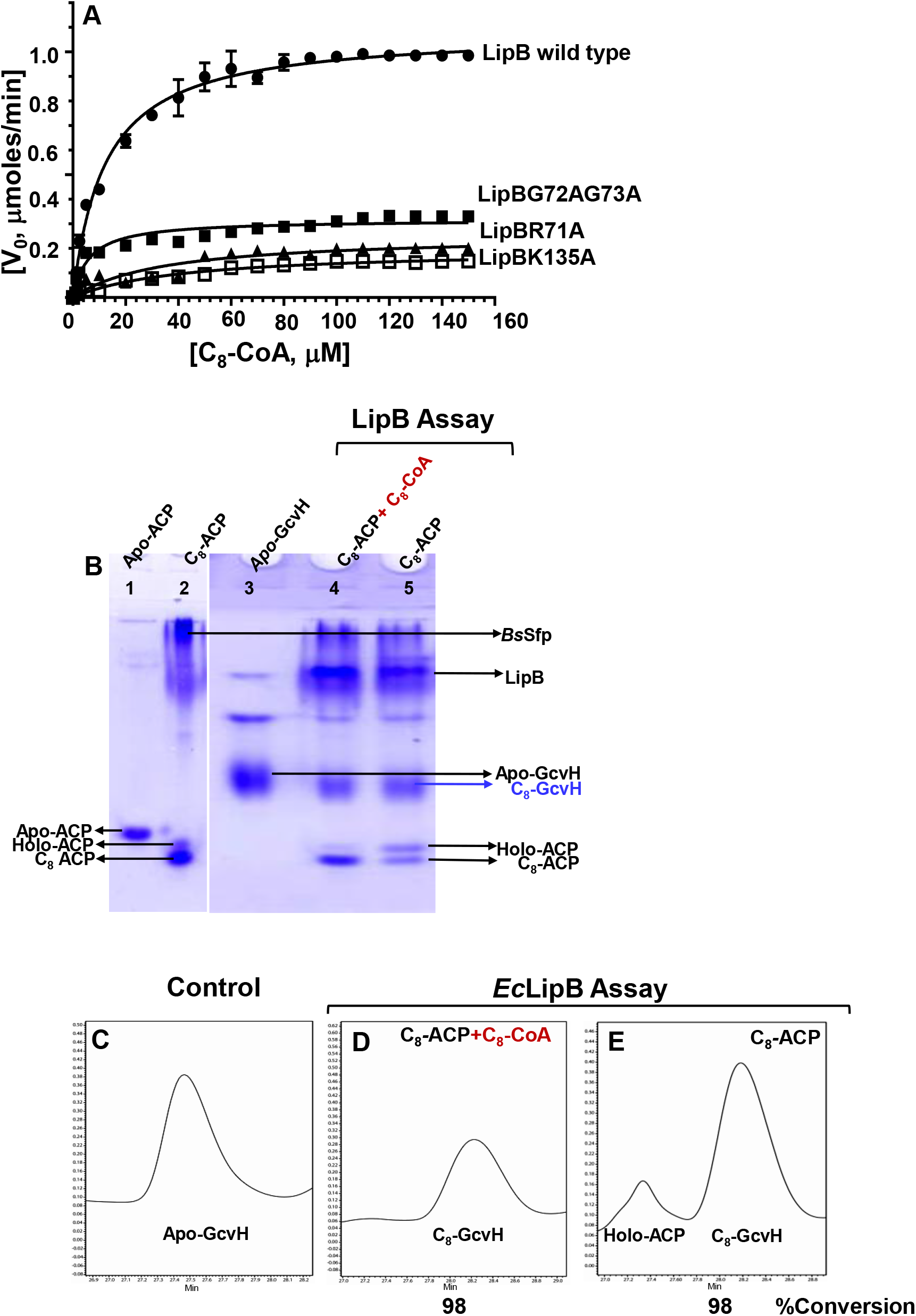
LipB prefers C_8_-CoA over C_8_-ACP as a donor substrate. A) Kinetic studies of GcvH conversion. using -●-LipB, -▴-LipBR71A, -▪-LipBG72A/G73A, and -□-LipBK135A in the presence of C_8_-CoA as a donor substrate and GcvH as an acceptor substrate. Reported values are the average of two independent measurements, done on a Symmetry C18 reversed phase HPLC column. The figure was drawn using GraphPad prism version 6.0. B)A 12% Native-PAGE gel, displaying Lane 1-3: apo-ACP, C_8_-ACP, and apo-GcvH, respectively used as control. Lane 4 was loaded with LipB assay performed in presence of equimolar concentrations of C_8_-ACP and C_8_-CoA used as donor substrates. Lane 5 shows the assay done using C_8_-ACP alone as a donor substrate. In both the assays, GcvH was used as an acceptor substrate for the bisubstrate reaction, and its conversion to C_8_-GcvH was followed. C18-reversed phase chromatogram for C) Apo-GcvH, LipB assay in presence of D) equimolar concentration of C_8_-ACP: C_8_-CoA, and E) C_8_-ACP alone. Samples in Fig. C, D and E, are equivalent to the samples in Lanes 3, 4 & 5 of the Native-PAGE gel in figure B.

### LipB can also octanylate GcvH using C_8_-CoA as a donor substrate

Next, we probed the importance of the ACP scaffold in substrate recognition by LipB. C_8_-CoA comprises a C_8_-chain and a phosphopantetheine arm, identical to the modifications present in C_8_-ACP. In addition, it contains an adenosine 3’5’-diphosphate group, not found in C_8_-ACP. Different length acyl-CoA were used as donor substrates *viz*. C_4_-, C_8_-, C_10_- and C_12_-CoA, and the reaction progress was monitored by following the modification of *E. coli* GcvH on an SDS-PAGE gel and MALDI-tof mass spectrometry. As shown in Fig. 2E, LipB efficiently modified apo-GcvH using C_8_-CoA as acyl-chain donor (lane 5). No change in GcvH band migration was observed when C_4_-, C_10_-, or C_12_-CoA were used as donor substrates, shown in Lanes 4, 6 & 7 in Fig. 2E. MALDI-tof mass spectrometry confirmed the increase in mass of GcvH by 127Da using C_8_-CoA as a substrate, equivalent to the molecular mass of the C_8_-chain (Fig. S4C). Figure S3A shows the mass spectra of apo-GcvH alone, and S3B-E mass spectra of GcvH after the LipB assay carried out using C_4_-, C_8_-, C_10_- and C_12_-CoA as acyl-chain donors. No change in mass was observed in the assays performed with C_4_-, C_10_- and C_12_-CoA.

Human LipT2 is an octanoyl transferase, associated with a vast number of disorders. Mice homolog (*Mus musculus*, UNP Q9D009) of LipT2 was used to check its ability to transfer octanoate from C_8_-CoA to *E. coli* GcvH. As shown in Fig. 2F, MmLipT2 was also able to use C_8_-CoA as a donor substrate. Mol. wt. marker was loaded in Lane 1, *E. coli* GcvH alone in lane 2, and *Mm*LipT2 assay product in lane 3.

Kinetic measurements were performed keeping LipB (5μM), GcvH (25μM) concentration constant, and varying the C_8_-CoA concentration (0-160 μM). The results were quantitatively analyzed on a C18-reversed phase column. Kinetic constants were obtained by performing hyperbolic Michaelis–Menten fits to the raw data, using GraphPad Prism version 6.0 (GraphPad Software, La Jolla, CA). A K_m_ value of 12.2 ±0.8 μM and a V_max_ of 1.1 ± μmolmin^-1,^ were obtained from the graph (filled circles), illustrated in Fig 3A. In a previous study using C_8_-ACP as a substrate, a K_m_ of 10.2±4.4μM has been reported for LipB(17).

### C_8_-CoA is preferred over C_8_-ACP as a LipB substrate

LipM, a *B. subtilis* homolog of LipB prefers C_8_-ACP over C_8_-CoA as a substrate (31). Thus, competitive assays were performed using equimolar concentrations of C_8_-ACP and C_8_-CoA, to understand the substrate preference of LipB. The conversion of C_8_-ACP and apo-GcvH into their holo-form was followed on a native-PAGE gel. Lanes 1, 2 and 3 in Fig. 3B display the bands corresponding to apo-ACP, C_8_-ACP, and apo-GcvH loaded as reference. Lane 4 was loaded with the assay performed in presence of equimolar concentration of C_8_-CoA and C_8_-ACP, and Lane 5 with the assay carried out using C_8_-ACP alone as a donor substrate. In lane 4, a major proportion of C_8_-ACP remained unused after the assay, though GcvH displayed complete conversion. On the contrary, an intense band equivalent to holo-ACP appeared in Lane 5, just above the C_8_-ACP band, in addition to the newly formed C_8_-GcvH band. C18-reversed phase chromatography confirmed the gel results. Fig. 3C shows a chromatogram for apo-GcvH alone, while Fig. 3D after the assay, displaying the peak for C_8_-GcvH formed during the assay. Fig. 3E displays the assay carried out using C_8_-ACP as a donor substrate. A distinct peak for holo-ACP appeared, formed by the removal octanoyl-chain from C_8_-ACP (Fig. 3E). This peak was not observed when the assay was performed in a reaction mixture containing equimolar C_8_-ACP and C_8_-CoA concentration (Fig 3D).

### LipB binds adenosine and phosphates of CoA

C_8_-CoA has an adenosine-3,5’-diphosphate group, in addition to the C_8_-chain and phosphopantetheine moiety. Octanoylation of GcvH by LipB using C_8_-CoA hints towards the presence of an adenylate binding site in LipB, akin to LplA. Thus, binding studies were performed using NMR to get insights into the atoms that interact. The chemical structure of Coenzyme A is shown in Fig. 4A, with all its protons labeled. Fig. 4B1 illustrates the ^1^H NMR spectra of Coenzyme A in buffer (50mM Tris, 200mM NaCl, pH 7.8). CoA chemical shifts were assigned based on the BMRB entry bmse000271. ^1^H Saturation Transfer Difference (STD) experiments were acquired using a Coenzyme A: LipB molar ratio of 8:1. Three peaks were observed in the 6.0-9.0 ppm range, corresponding to 2, 8 and 1’ resonances of adenosine 3’5’-diphosphate component of CoA. Fig. 4B2 illustrates the STD spectra obtained for CoA in presence of LipBK135A/C169A (catalytically inactive double mutant). NOE peaks corresponding to position 1’ and 8 were observed, suggesting interaction of the adenine ring of CoA with LipBK135A/C169A. In all the experiments, 8:1 CoA: LipB molar ratio was used.

**Fig. 4.**
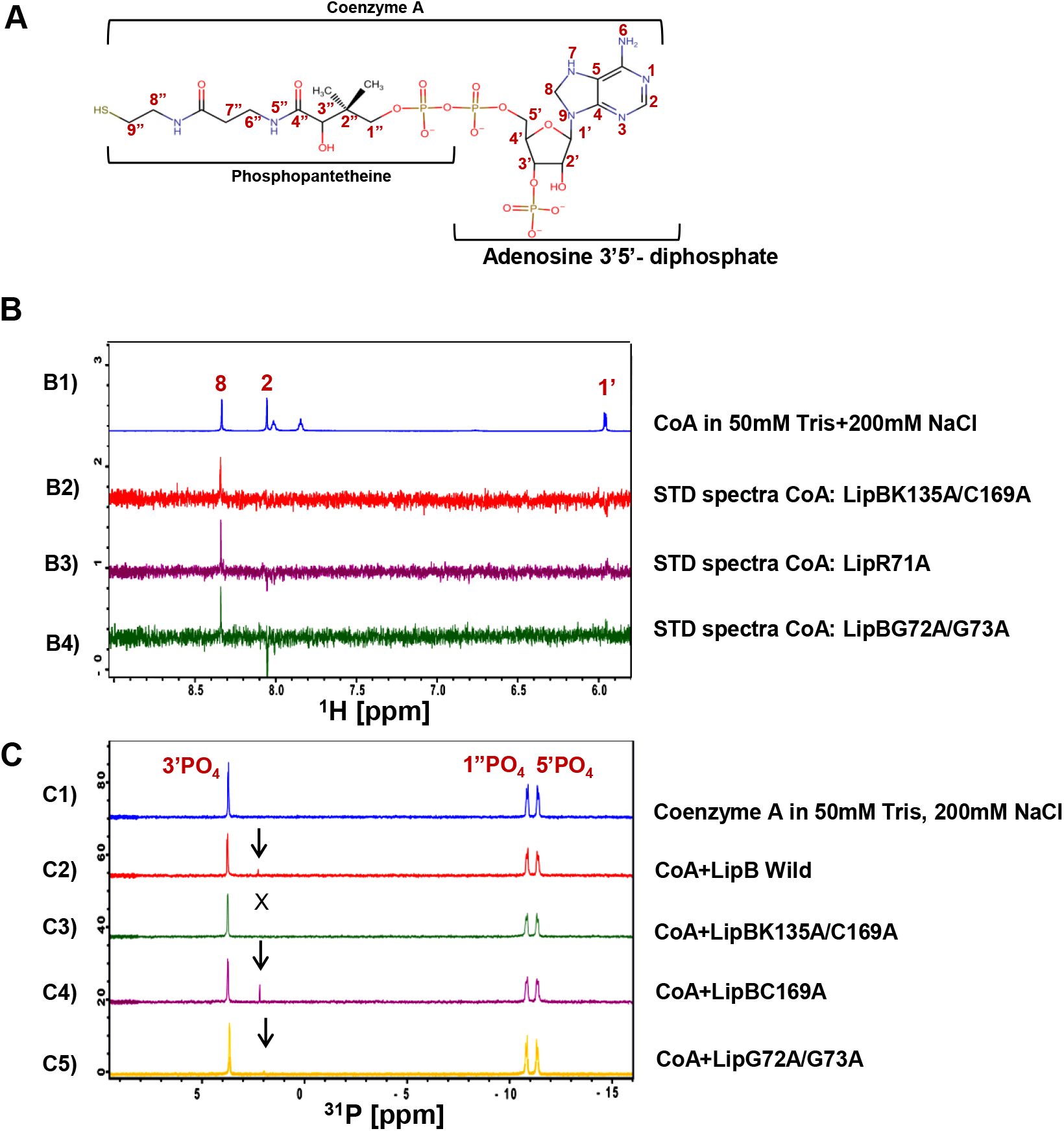
LipB interacts with adenosine 3’5’-diphosphate of CoA. A) The chemical structure of Coenzyme A is shown, displaying its components Phosphopantetheine and ADP. B1) ^1^H NMR spectra of CoA in buffer, B2) Saturation transfer difference (STD) spectra for CoA: LipBK135A/C169A, B3) STD spectra for CoA: LipBR71A B4) STD spectra for CoA: LipBG72A/G73A. C) ^31^P NMR spectra of C1) CoA in buffer, C2) CoA: Wild type LipB, C3) CoA: LipBK135A/C169A, C4) CoA: LipBC169A, C5) CoA:LipBG72A/G73A. The molar ratio of CoA: LipB was maintained at 8:1 in all the experiments.

^31^P NMR studies were also performed to verify the interaction of LipB with phosphates of CoA, if any. Fig. 4C1 shows the ^31^P NMR spectra of CoA in 50mM Tris, 200mM NaCl, pH 7.8. Three peaks were observed, corresponding to 3’-, 1”-and 5’-phosphate. Fig. 4C2-C4 show the spectra for CoA in presence of wild-type LipB, LipBK135A/C169A and LipBC169A, respectively. The peak corresponding to 3’-phosphate displayed an upfield change in chemical shift upon addition of wild type LipB (Fig. 4C2) and LipBC169A (Fig. 4C4). No change in peak position was observed in the double mutant LipBK135A/C169A (Fig. 4C3), suggesting direct interaction of Lys135 with 3’-phosphate of CoA. The new peak arising as a result of this interaction has been indicated by a downward pointing arrow in the figures. A CoA:LipB ratio of 8:1 was used for 31P NMR studies.

### Adenine and phosphate binding sites are conserved in *Mt*LipB structure

Structural insights into the CoA-LipB interaction were obtained by superimposing *M. tuberculosis Mt*LipB structure (PDB 1W66) on the catalytic domain of *E. coli* Lipoate-protein ligase A complexed with C_8_-AMP (LplA, PDB 3A7A), shown in Fig. 5A (32). In most proteins, atoms N2, N6, C2 and C8 of adenine, present at the Watson-Crick edge interact with the backbone and side chain atoms of adenine binding proteins (33). In LplA (PDB 3A7A), N1 and N6 of adenine form a hydrogen bond with the side chain of Asn 83 OD1, Thr 151 and the backbone O of Phe 78. Also, N7 forms a hydrogen bond with the backbone N of Phe 78. In *M. tuberculosis* LipB (*Mt*LipB, 1W66), remarkably similar residues are present at these sites, instead of Asn 83 and Phe 78, Gln 87 and Trp 82 are present. These residues correspond to Gln 82 and Tyr 77 in *E. coli* LipB, while in *Mm*LipT2, Gln 96, and Phe 91 are present. The oxygen atoms of AMP (O4 and O5) interact with the ε amino group of Lys 133 in LplA. The equivalent residue is Lys 142 in *Mt*LipB, and Lys 135 in *E. coli* LipB. Also, in the LplA structure (PDB 3A7A), lipoyl moiety interacts with numerous hydrophobic/aromatic side chains, *viz*. Trp 37, Val 44, Arg 70, Val77, His79, and His149. In *Mt*LipB, these positions are occupied by Leu 47, Thr 54, Arg 76, Thr 81, His 83, and His 157 (Fig. 5B). In *E. coli* LipB, Val 41, Thr 48, Arg 71, Thr 76, His 78, and His 150 are present at the same position in the sequence, shown in Fig. 4B.

**Fig. 5.**
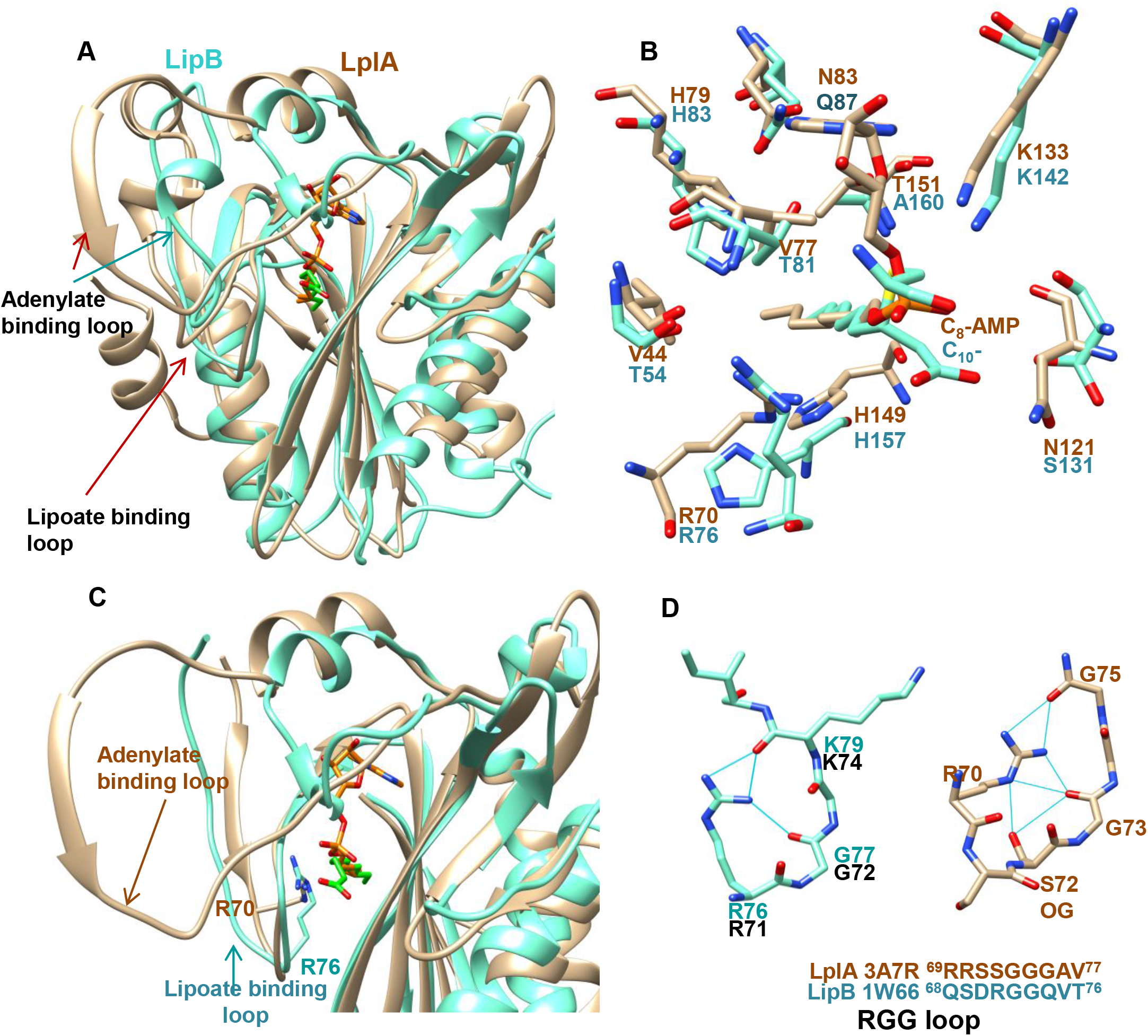
*M. tuberculosis* LipB has an adenine and phosphate binding site conserved within its active site cavity. A) Ribbon representation of *M. tuberculosis* LipB (PDB 1W66, colored cyan) overlaid on the *E. coli* LplA catalytic domain structure (PDB 3A7R, colored brown). B) LplA side chains that interact with lipoyl-5-AMP are colored light brown. The corresponding residues in *M. tuberculosis* LipB (PDB1W66) are colored cyan. C) The lipoate binding loop of LplA overlaid on the RGG loop of *Mt*LipB. D) Side chain to backbone hydrogen bond interactions of Arg76 with the backbone carbonyls of residues present in the RGG loop of *Mt*LipB (1W66) are shown. These interactions are similar to the side chain interactions of Arg (3A7R).

The lipoate binding loop ^69^RRXXGGG^75^ of LplA (PDB 3A7A) is also partially conserved in *Mt*LipB, and has a very similar conformation, shown in Fig. 5C and D (34). MtLipB backbone and side chain are colored cyan, while LplA is colored brown. Arg 70 side chain and Gly 73 backbone of LplA structurally overlap with Arg 76 and Gly 78 of *M. tuberculosis* LipB. Arg 71 and Gly 72 are the equivalent residues in *E. coli* LipB. Interestingly, Arg 70 side chain forms unique side chain to backbone hydrogen bonds with the carbonyls of Gly73, Gly75 and side chain of Ser72 in the LplA structure (Fig. 5D, colored brown). These interactions have been speculated to stabilize the lipoate binding loop of LplA. In *Mt*LipB as well, arginine 76 side chain forms identical hydrogen bonds with the backbone carbonyls of Gly 77 and Lys 79, colored cyan shown in Fig. 5D.

Thus, the structural comparison of *E. coli* LplA crystal structure with *M. tuberculosis* LipB structure suggests conservation of an adenylate binding site, comparable to LplA.

### An ‘RGG’ loop is necessary for LipB catalytic activity

RGG loop is structurally similar to the lipoate binding loop of LplA. The importance of this loop in LipB function remains unclear. Thus, LipBR71A and LipBG72A/G73A mutants were generated and assays performed using C_8_-CoA and GcvH as donor and acceptor substrates. Fig. 6A shows an SDS-PAGE gel, displaying a molecular weight marker in Lane 1, purified apo-*Ec*GcvH in Lane 2, and assays carried out with wild type LipB, LipBG72A/G73A, LipBR71A, LipBK135A, LipBK135A/C169A, respectively in lanes 2-6, using C_8_-CoA as an octanoate-donor and GcvH as octanoate acceptor. Apo-to C_8_-GcvH conversion was observed based on the forward migration of the GcvH band in the gel. Compared to wild type LipB, conversion was remarkably reduced in Lanes 3, 4, 5, and 6, suggesting decreased activity of the mutants. LipBG72A/G73A displayed 39% while LipBR71A displayed 21% activity compared to wild type, determined using C18 HPLC reversed phase chromatography.

**Fig. 6.**
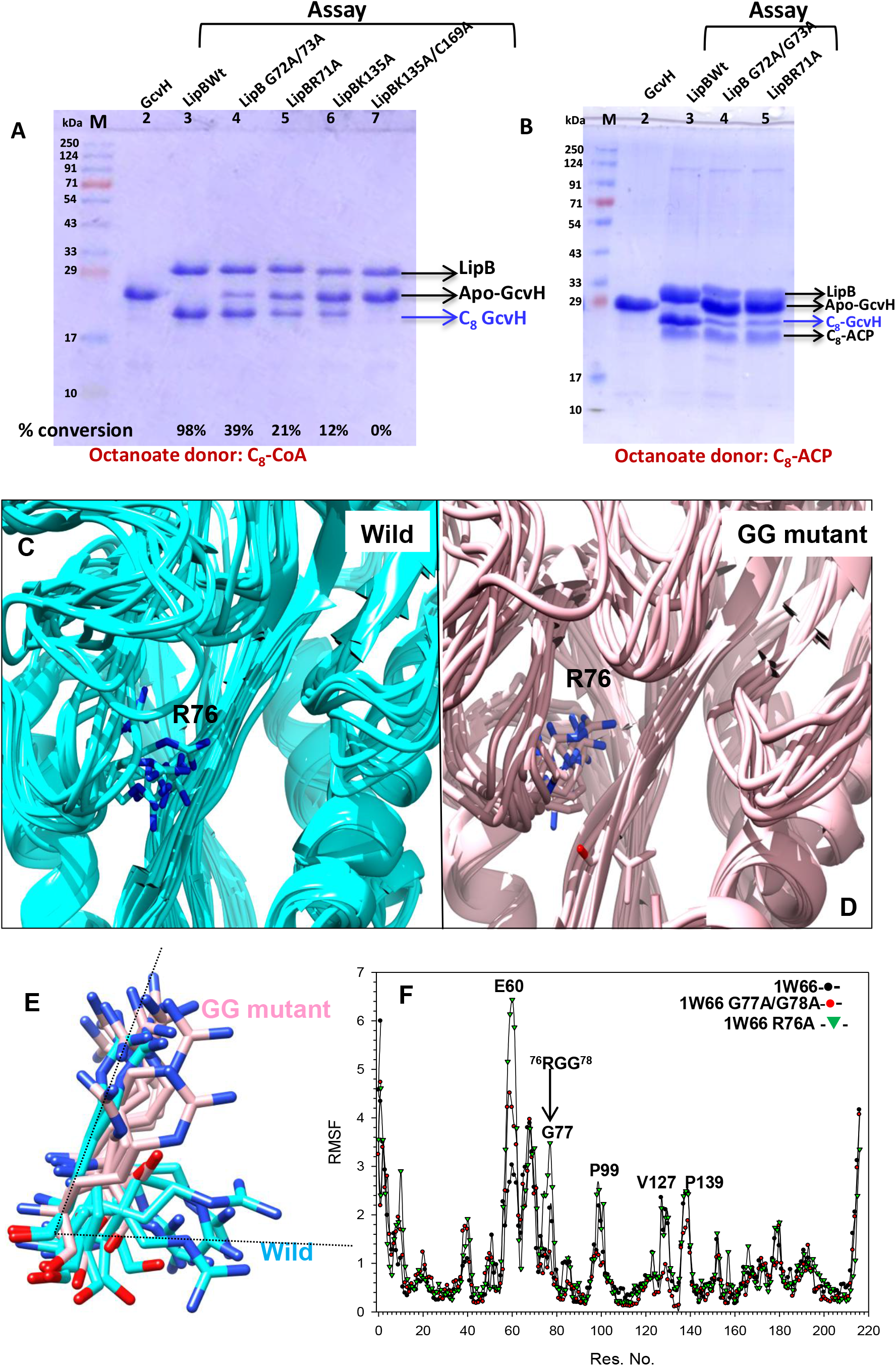
The RGG loop is necessary for LipB function. A) A 12% SDS-PAGE gel, displaying molecular weight marker in Lane 1, Lane 2 is GcvH alone, Lanes 3-7; assay carried out in presence of wild type LipB, LipBG72A/G73A, LipBR71A, LipBK135A, and LipBK135A/C169A, respectively, using C_8_-CoA as an octanoate-donor. Percent conversion of GcvH after the assay is shown at the bottom of each lane. B) A 12% SDS-PAGE gel, displaying Lane 1, apo-GcvH, Lane 2-4; assay carried out with wild type LipB, LipBG72A/G73A and LipBR71A, using C_8_-ACP as a donor substrate. C) Lowest 10 models generated for *Mt*LipB wild type (cyan) and D) *Mt*LipBG77A/G78A mutant (pink).E) Arg 76 side chain conformation in the 10 lowest energy models generated for *Mt*LipB wild type (cyan) and *Mt*LipBG77A/G78A mutant (pink). F) Root mean square fluctuation profiles for wild type *Mt*LipB (black filled circle), *Mt*LipBG77A/G78A (red filled circle) and *Mt*LipBR76A (green triangles). RMSF values were derived from simulation trajectories obtained from CABS-flex 2.0.

Similar LipB assays were also performed using C_8_-ACP as octanoate-donor substrate and GcvH as acceptor. A marked reduction in enzyme activity was observed in LipBG72A/G73A and LipBR71A mutants, compared to wild type (Fig. 5B). In the gel, Lane 1 is molecular weight marker, lane 2 is apo-GcvH (without assay); Lanes 3-5 GcvH after the assay with wild type LipB, LipBG72/G73A and LipBR71A mutants, respectively. Kinetic measurements were also performed on the LipBR71A (filled triangles) and LipBG72A/G73A mutants (filled squares), using C_8_-CoA as an octanoate-donor (Fig. 3A). A K_m_ value of 6.7 ±0.9μM (a 2-fold decrease) and a V_max_ of 0.32 ± 0.01 μmolmin^-1^ (∼3-fold decrease compared to wild type) was observed for the LipBG72A/G73A mutant. LipBR71A mutant displayed a K_m_ value of 34.4 ±8.8 μM (∼3fold increase) and a V_max_ of 0.26± μmolmin^-1^ (4-fold decrease, Fig. 3A). In comparison, LipBK135A mutant displayed a K_m_ of 42.4 ±4.6 μM (3.5 fold change) and a V_max_ of 0.2±0.01μmolmin^-1^.

Saturation transfer difference experiments confirmed the binding of CoA with LipBG72/G73A and LipBR71A mutants equivalent to wild type LipB. Fig. 4B3 and Fig 4B4 display 1D Saturation Transfer Difference (STD) spectra for CoA: LipBR71A and CoA: LipBG72A/G73A interaction. NOEs matching positions 2, 8 and 1’ of adenine ring were observed. Figure 4C5 shows the ^31^P NMR spectra for the interaction of CoA with LipBG72A/G73A mutant. The 3’-phosphate peak displayed a chemical shift change similar to wild type protein, suggesting phosphate binding.

### Molecular Dynamics simulations suggest that the RGG sequence imparts conformational flexibility to the loop

Notably, the RGG loop forms one side of active site cavity. Therefore, molecular dynamics simulations were performed on PDB 1W66 (*Mt*LipB structure) using CABS flex 2.0 to identify motions of the RGG loop(35). PDB 1W66 was edited to generate LipBR76A and LipBG77A/G78A structures. Figure 6C & D display the conformation of the ‘RGG’ loop in the 10 models generated by MD simulations, using PDB 1W66 for wild type and the *Mt*LipBG77AG78A mutant. In wild type *Mt*LipB, Arg 76 side chain appears to flip out of the active site cavity in 8 out of 10 models (Fig. 6E), while in LipBG77A/G78A mutant all Arg 76 side chains faced the cavity in the 10 generated models, suggesting restrained side chain motions in the double glycine mutant. Figure 6F displays RMSF values for the simulations as a function of residue number for wild type *Mt*LipB (PDB 1W66, filled black symbols), *Mt*LipBR76A (inverted green) and *Mt*LipBG77A/G78A mutant (filled red). In *Mt*LipBG77A/G78A mutant, rmsf values decreased in magnitude, while in the *Mt*LipBR76A mutant, rmsf values increased drastically for the ‘RGG’ loop compared to wild type. A loop in the capping sheet also displayed increased rmsf values in both the mutants compared to wild type. Overall, the results suggest that the mutations cause a noticeable change in dynamics of the RGG loop, as well as the nearby loops present in the capping sheet.

## DISCUSSION

In prokaryotes, two different pathways for lipoic acid modification coexist, a) an endogenous lipoic acid synthesis pathway, catalyzed by the consecutive action of Lipoate protein ligase B (LipB) and Lipoyl synthase (LipA), or b) a salvage pathway carried out by Lipoate protein ligase A (LplA). Yeast and higher eukaryotes lack the scavenging pathway, and derive lipoic acids exclusively from the biosynthetic pathway by the successive action of LipT2, LIAS and LipT1, metazoan homologs of LipB, LipA and LplA, respectively(5). The three enzymes work in concert, performing non-redundant functions in the mitochondria. Genetic mutation of any one, or the upstream type II fatty acid synthesis enzymes causes metabolic dysfunction (36,37). The disorders due to malfunction of lipoic acid synthesis pathway cannot be treated by supplementation with exogenous lipoic acid, as none of the three enzymes can uptake free lipoic acid. Thus, there is pressing need to better understand lipoic acid synthesis at the molecular level. (11, 21, 22).

LipB differs from the salvage enzyme LplA in several respects; a) C_8_-ACP, an intermediate of fatty acid synthesis pathway is the only known substrate of LipB, whereas LplA is a lipoic acid/ATP dependent enzyme. b) LipB uses cysteine-lysine dyad as catalytic residues, while a lysine performs lipoate adenylation as well as transfer reactions of LplA (34). c) The catalytic reaction proceeds *via* different intermediates; LplA forms a non-covalent lipoyl-5’-AMP intermediate from lipoic acid and ATP, while a C_8_-LipB covalent adduct is formed using mitochondrial fatty acid synthase intermediate (C_8_-ACP). Despite these dissimilarities, unique parallels also exist between the two enzymes with regard to substrate recognition. Saturation transfer difference experiments suggest conservation of an adenine binding site in LipB, similar to LplA. Likewise, our ^31^P NMR studies disclose an interaction between 3’-phosphate of adenosine and Lys 135, similar to Lys 133 of LplA (38). Stronger affinity of holo-ACP for LipB, and perturbations observed in one of the NH resonances of the phosphopantetheine moiety in HSQC titration experiments also substantiate the presence of a phosphate binding site in LipB. Structural overlay of *Mt*LipB (1W66) with the catalytic domain of *E. coli* LplA (3A7A) suggests conservation of residues in LipB at sites that interact with the adenylate moiety and phosphates of AMP in LplA structures (PDB 3A7A, 2ART, 2ARU). Interestingly, the two sites are conserved not just in LipB but also Lipoate transferase 2 (LipT2, UNP A6NK58) as reported by the assay using C_8_-CoA, divulging evolutionary relationships(39).

The salvage enzyme LplA interacts with lipoic acid by means of a well-defined lipoate-binding loop. An ^76^RGG^78^ loop present in the active site of (*M. tuberculosis*) *Mt*LipB (1W66) has a similar sequence and conformation compared to the lipoate binding loop ^69^RRSSGGGAV^77^ of LplA. In the LplA structure (PDB 3A7A), Arg 69 forms side chain to backbone hydrogen bonds with the glycine and Ala of the loop, in a manner similar to the Arg76 of LipB. The lipoate binding loop is crucial for lipoylation in LplA, and experiences large scale conformational changes upon lipoic acid binding(34). MD simulations of *M. tuberculosis* LipB (PDB 1W66) suggest comparable motions of the RGG loop in LipB as well, that are quenched upon mutation of the two glycine’s in the loop to alanine. Conversely, mutation of the conserved Arg in the RGG sequence to an Ala increased the overall dynamics of the loop, inducing major changes in conformation of the active site. ‘RGG’ sequences are frequently observed in RRM domains, and are known to bind phosphates of nucleotides/quadruplexes(40). Electrostatic attraction/hydrogen bonds between positively charged guanidinium of arginine and the negatively charged phosphates contribute to this interaction(41,42). Interactions between phosphate and Phosphotransacetylase enzyme from *Methanosarcina thermophila* have also been attributed to arginines(43). We surmise, that the ‘RGG’ loop of LipB is necessary for maintaining flexibility of the active site cavity, and may also be transiently involved in interaction with phosphates.

Taken together, the present study highlights the importance of ACP modifications in LipB recognition. Apo-ACP weakly binds LipB, while the presence of a phosphopantetheine moiety (in holo-ACP) allows ACP to bind with several-fold higher affinity. Addition of a C_8_-chain further increases binding affinity. These observations are in sync with two recent papers that show LipB can recognize different acyl intermediates allosterically, without even flipping the acyl chain (29,30). The dissociation constant obtained for C_8_-ACP:C169ALipB interaction in one of those studies using NMR (8.0±2.64μM), is in close agreement with the values we obtained using SPR (5.5± 0.03μM) (30). The pattern and magnitude of chemical shift change is also consistent with our studies (30). Bartholomew et al observed a more stable complex between C_8_-ACP and LipB, compared to other acyl-ACP’s using MD simulations, which was attributed to higher surface complementarity of the two proteins. Contribution of the phosphopantetheine arm (4’-PP) in ACP recognition by LipB was however not taken into consideration. In the MD generated structures of the intermediates, 4’-phosphopantetheine arm conformation is remarkably different (ref#29, Supplementary Fig. S10B therein). This difference in conformation of the 4’-PP arm was also echoed by the chemical shift of Ser 36 (covalently attached to 4’-PP), that displayed a noticeable change in magnitude in the different intermediates, *viz*. C_6_-, C_8_- and C_10_-ACP upon LipB titration (ref#29). Comparison of the *E. coli* C_6_-(PDB 2FAC) and C_10_-ACP (PDB 2FAE) crystal structure also provides similar insights into the difference in conformation of phosphopantetheine moiety. Thus, recently published studies, as well as our own support the hypothesis that 4’PP arm contributes significantly to ACP-LipB interaction.

Above all, our studies disclose a new substrate analog of C_8_-ACP, *i*.*e*. C_8_-CoA, that can serve as a substrate for LipB, as well as LipT2. Though C8-CoA concentration is tightly regulated in the mitochondria, this information might be of assistance while treating patients with human mitochondrial fatty acid synthase mutations; genes encoding (MECR) mitochondrial trans-2-enoyl-coenzyme A-reductase(21,44), malonyl-CoA synthetase (ACSF3), malonyl CoA-acyl carrier protein transacylase (MCAT) *etc*. characterized by neurological manifestations due to the loss of lipoylation(21). The symptoms overlap with CoPAN and PKAN disorders (defects in CoA biosynthesis), as all these conditions have the same consequence, *i*.*e*., reduction in lipoylation (26). Two candidates have been proposed as a therapy so far; lipoic acid and octanoic acid. Studies with radioactive lipoic acid supplements show that exogenous lipoic acid enters the blood stream and tissues, but is rapidly degraded by β-oxidation, with no noticeable lipoic acid modification (42). Likewise, in LipT2, LIAS or LipT1 knockouts, lipoic acid supplements do not provide any respite(5). We surmise that octanoic acid analogs might serve as a better substitute for C_8_-ACP, due to their ability to pass through the mitochondrial membrane, convert into octanoyl-CoA inside the mitochondria, and act as a donor substrate for LipT2. Support for this proposition also comes from studies on individuals with medium-chain acyl-CoA dehydrogenase deficiency (MCADD), a fatty acid oxidation (FAO) defect, characterized by elevated C_8_-carnitine concentration, and lack of acyl-CoAs >C10CoA. These individuals display increased lipoylated pyruvate dehydrogenase complex (PDC) levels compared to healthy controls, associating medium chain CoA to lipoic acid synthesis(45). Another recent study on yeast with deleted 3-hydroxyacyl thioester dehydrogenase, an important enzyme of mitochondrial FAS (Δhtd2), but harbouring a mitochondrially mislocalized fatty acyl-CoA ligase (Fam1-1 suppressor allele), was able to rescue the cells by growth on C_8_-supplemented media. FAM1-1 has been long back shown to activate fatty acids by attachment to CoA or ACP (46). Authors attributed the suppression of respiratory growth defects in cells lacking mtFAS enzymes by Fam1-1, to a LipB dependent mechanism, as this suppression could not be achieved in the absence of lipoylating enzymes or C_8_-supplements (47). Thus, our *in vitro* studies validate their findings, offering proof of concept that LipB/LipT2 can indeed efficiently use C_8_-CoA as a substrate, apart from C_8_-ACP. Further studies are required, to test the efficacy of octanoate-in rescuing FAS deficient mutants.

## Materials and Methods

### Cloning, Expression, and Purification

Genes encoding ACP, LipB, and GcvH were PCR amplified from *E. coli* genome, and cloned in a pET28a expression vector. Mice LipT2 gene (synthesized after codon optimization) was also cloned similarly. Protein expression was induced with isopropyl β-D-1-thiogalactopyranoside, and purified by Ni^2+^-NTA chromatography, followed by size exclusion chromatography. Apo-ACP was converted to Holo-ACP and C_8_-ACP *in vitro*, using *Bacillus subtilis* phosphopantetheinyl transferase (Sfp), using a modified protocol of Lambalot and Walsh (48). Sfp was expressed and purified by ion exchange, followed by Size exclusion chromatography. Uniformly labeled [^1^H,^15^N,^13^C] holo-ACP and C_8_-ACP were prepared by growing *E. coli* Bl21(DE3) cells in M9 medium containing ^15^N NH_4_Cl (1g/L) and ^13^C glucose (2g/L). For NMR measurements, N-terminal His tag of ACP was removed by thrombin cleavage, using immobilized thrombin. Mutagenesis studies were performed using site-directed mutagenesis approach, and purified using a protocol similar to that of the wild-type protein.

### NMR Data Acquisition

NMR samples comprised uniformly labeled [^1^H,^15^N,^13^C] protein, in 50mM Tris-HCl, 200mM NaCl, pH 8.0, 90% H_2_O, and 10% D_2_O, 0.5% sodium azide. Two-dimensional NMR experiments, *viz*. ^1^H^15^N HSQC, ^1^H^15^N TROSY were acquired on a Bruker Avance III 700 MHz NMR spectrometer, installed at the National Institute of Immunology (New Delhi, India), equipped with triple resonance TXI probe. ^1^H^15^N TROSY-HSQC spectra were acquired using 1024 data points in the *t*_2_ dimension and 512 data points in the *t*_1_ dimension. Saturation transfer difference experiments were acquired using Bruker pulse sequence. The ligand peaks observed in the 1D STD experiment were assigned based on ^1^H NMR spectra of CoA. 1D ^31^P measurements were done on a BBI (broad band inverse) probe, and the experiments were performed for ∼8h. NMR data were processed on a workstation with Red Hat Enterprize Linux 5.0, using NMRDraw/NMRPipe, and analyzed using Sparky(49). Experiments were performed at 298 K throughout. ^15^N^13^C spectra were indirectly referenced using a chemical shift standard, sodium 2,2-dimethyl-2-silapentane-5-sulfonate (DSS) (45).

### NMR Data Analysis

Chemical Shift Perturbations (CSP) have been reported as weighted average of the nitrogen and ^1^H chemical shifts (ΔAvg_HN_), derived from equation-1(50)

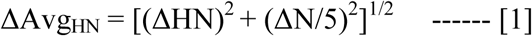

where ΔHN and ΔN are the changes in the proton and nitrogen dimensions, respectively. One standard deviation has been used as a cutoff to mark significant change.

### NMR titration studies

^15^N labeled C_8_-, apo- or holo-ACP in 50mM Tris, 200mM NaCl, pH 7.8, was titrated with LipBK135AC169A, and chemical shift perturbations were followed in a ^15^N TROSY-HSQC experiment. ^15^N labeled Holo- and octanoyl-ACP were first prepared by converting apo-ACP to C_8_-ACP using *Bacillus subtilis* phosphopantetheinyl transferase. For each titration point ^1^H^15^N TROSY spectra was acquired.

### Binding Studies using SPR

Binding strength of LipB for its substrates was determined by Autolab ESPRIT Surface Plasmon Resonance spectrometer, and analyzed by Autolab SPR Kinetic Evaluation Software. Gold surface was activated by N-hydroxysuccinimide (NHS, 0.05 M)/N-ethyl-N-(diethylaminopropyl) and carboiimide (0.2 M). LipB was immobilized on the gold surface in two channels. Unoccupied sites were blocked with 100 mM ethanolamine. After each experiment, regeneration was carried out with 50 mM NaOH. Increasing concentration of the substrate was injected on the sensor surface. Association and dissociation kinetics were monitored for 300s and 150s, respectively in 20 mM sodium phosphate buffer pH 8.0, and 100 mM NaCl. All experiments were carried out at 298K.

### Enzyme Assay

Enzyme activity was performed using 5 µM LipB, 75 µM C_8_-CoA/octanoyl-ACP as the donor substrate, and 25 µM GcvH as the acceptor substrate in 50mM Tris-HCl, 200 mM NaCl, pH 8.0, at 30°C. After 2 hr incubation, samples were loaded on a 12 % SDS-PAGE to follow GcvH conversion, or Native-PAGE gel to follow breakdown of C_8_-ACP. For quantitative measurements, samples were loaded on a Symmetry C18 5μm reversed phase column, on a Waters HPLC. Integrals of the area under the C_8_-GcvH curve were used as a measure of enzymatic conversion. For LipT2 assay, the reaction mixture was incubated at 37°C, and the buffer comprised 50mM Tris-HCl, 500 mM NaCl, pH 8.0. The octanoate-acceptor substrate was GcvH in all the assays.

### Mass Spectrometry

The assay samples were passed through C_4_ Zip Tips to remove salt, and the final sample eluted in 50 % acetonitrile and 0.1 % trifluoroacetic acid for analysis in Applied Biosystems 4800 MALDI-TOF spectrometer.

### CABS flex simulations

Coarse-grained simulations (10ns length) of protein motions were done using 50 cycles, and 50 cycles between trajectory frames, and the 10 final generated models and their trajectories were analyzed (35).

## Abbreviations used

LipB: Lipoate protein ligase B
ACP: Acyl carrier protein
GcvH: Glycine cleavage system H protein

## Contributions

U.Y., C.D., S.K. and M.S. conceived the project, designed the experiments, and interpreted the results. M.S. wrote the paper, with help from C.D. and U.Y.

## Acknowledgements

Authors thank the Department of Biotechnology, Govt. of India, and SERB (Science and Engineering Research Board) Sanction order No. “EMR/2017/002093” for financial support. Fellowship to CD from the Lady Tata Foundation, India, and UY from the University Grants Commission, India is gratefully acknowledged. Authors also thank Ms. Shanta Sen, Mass Spectrometry Facility, NII, for help with Mass Spectrometry data, and the Advanced Instrumentation Research Facility, Jawaharlal Nehru University, India for SPR measurements.

## Figure Legends

**Table-1.** Enzyme kinetic parameters for wild type LipB and the mutants.

|  | LipBWt | LipBG72A/G73A | LipBR71A | LipBK135A |
| --- | --- | --- | --- | --- |
| $V_{\max}$ ($\mu\text{moles/min}$) | $1.08 \pm 0.01$ | $0.32 \pm 0.01$ | $0.26 \pm 0.02$ | $0.20 \pm 0.01$ |
| $K_m$ ($\mu\text{M}$) | $12.24 \pm 0.79$ | $6.71 \pm 0.98$ | $34.39 \pm 8.8$ | $42.37 \pm 4.58$ |
| $k_{\text{cat}}$ ($\text{min}^{-1}$) | 0.22 | 0.06 | 0.05 | 0.04 |

The assay was performed using $\text{C}_8$ -CoA as the octanoate donor and *E. coli* GcvH as octanoate acceptor.

