## Supplementary material for "Lipoate protein ligase B primarily recognizes the C_8_-phosphopantetheine arm of its donor substrate, and weakly binds the acyl carrier protein": Allsupplementary.pdf

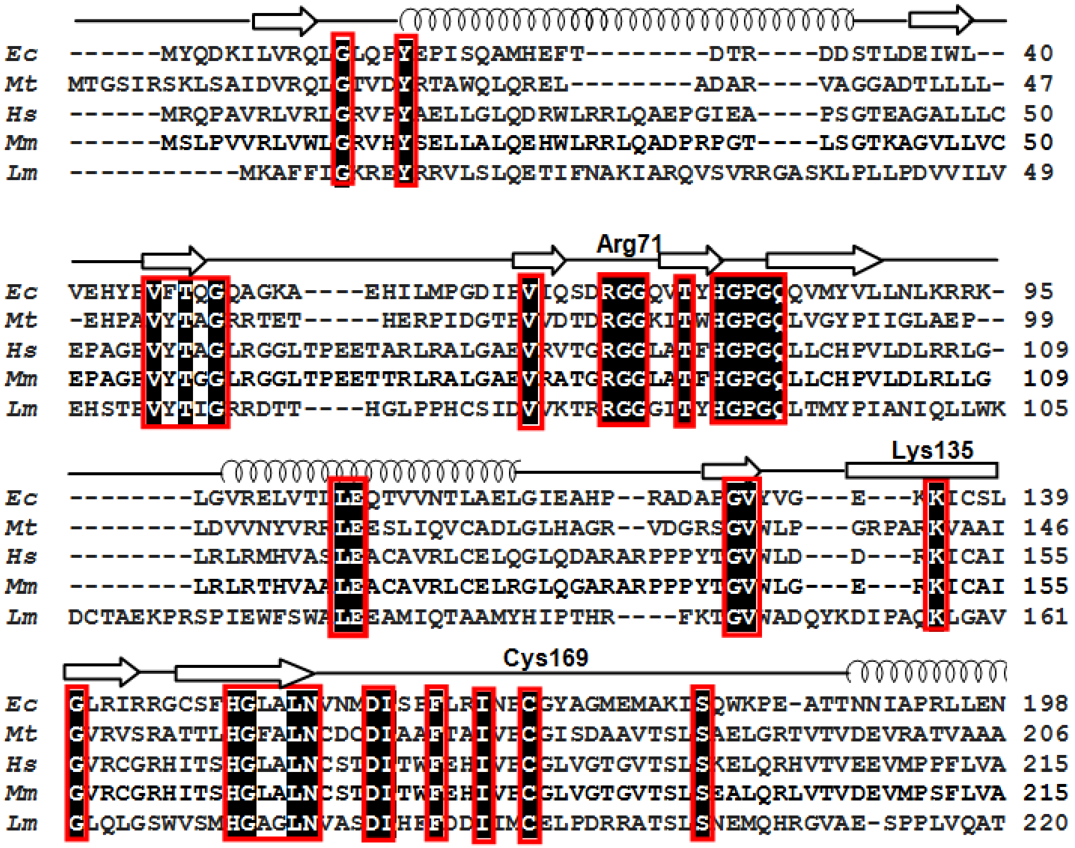


**Fig. S1** Sequence comparison of *E. coli* LipB with octanoyl-transferases from other sources. Conserved residues are shown as red boxes.

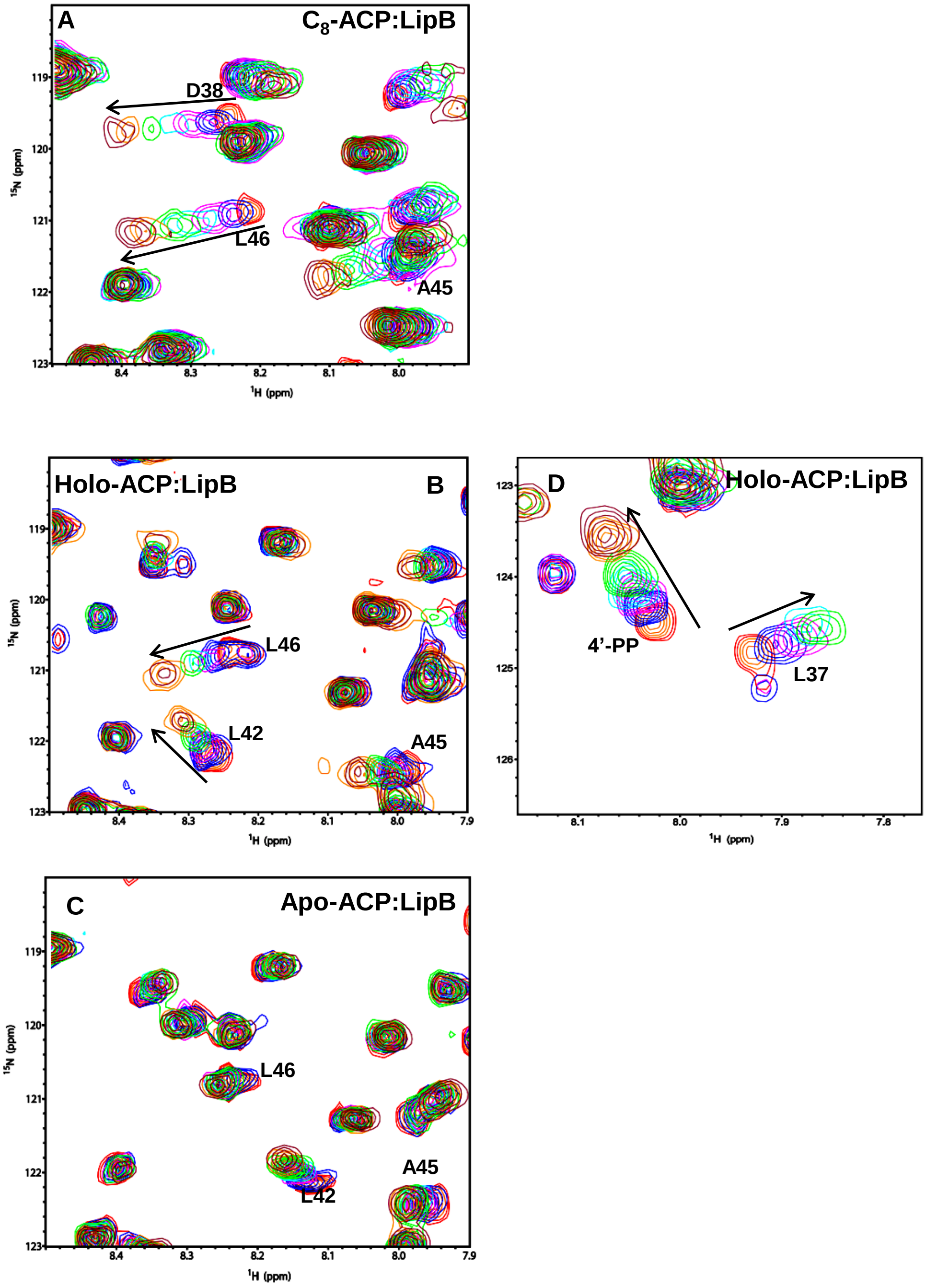


**Fig. S2.** Multiple overlaid $^{1}H^{15}N$ TROSY-HSQC spectra for the titration of ACP with LipBK135A/C169A. A region of the $^{1}H^{15}N$ TROSY-HSQC spectra for A) $C_8$-ACP, B) holo-ACP, and C) apo-ACP, upon titration with LipBK135AC169A. Red peaks represent free ACP, blue: 1:0.25, magenta 1:0.5, cyan 1:0,75, green 1:1, orange 1:1.5 and maroon 1:2 ACP: LipBK135A/C169A molar ratio. Residues that display significant perturbations are labeled.

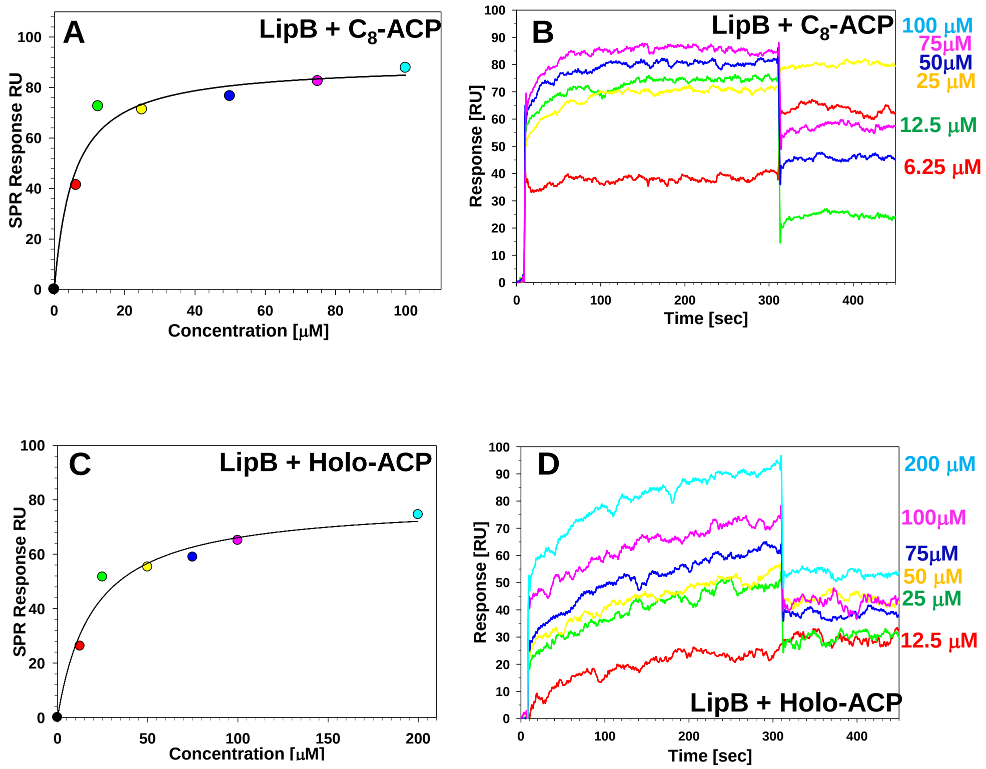


**Fig. S3.** Surface Plasmon Resonance measurements for the interaction of LipB with ACP. SPR sensorgrams for the binding of immobilized LipBK135A/C169A with A) $C_8$-ACP and C) holo-ACP. The color of different sensorgrams correspond to the concentration mentioned on the right side of the figure, that was used to pass over the immobilized sample. The plot of maximum response reached at equilibrium (Req) for each concentration of ligand used in the sensorgrams for B) $C_8$-ACP and D) holo-ACP.

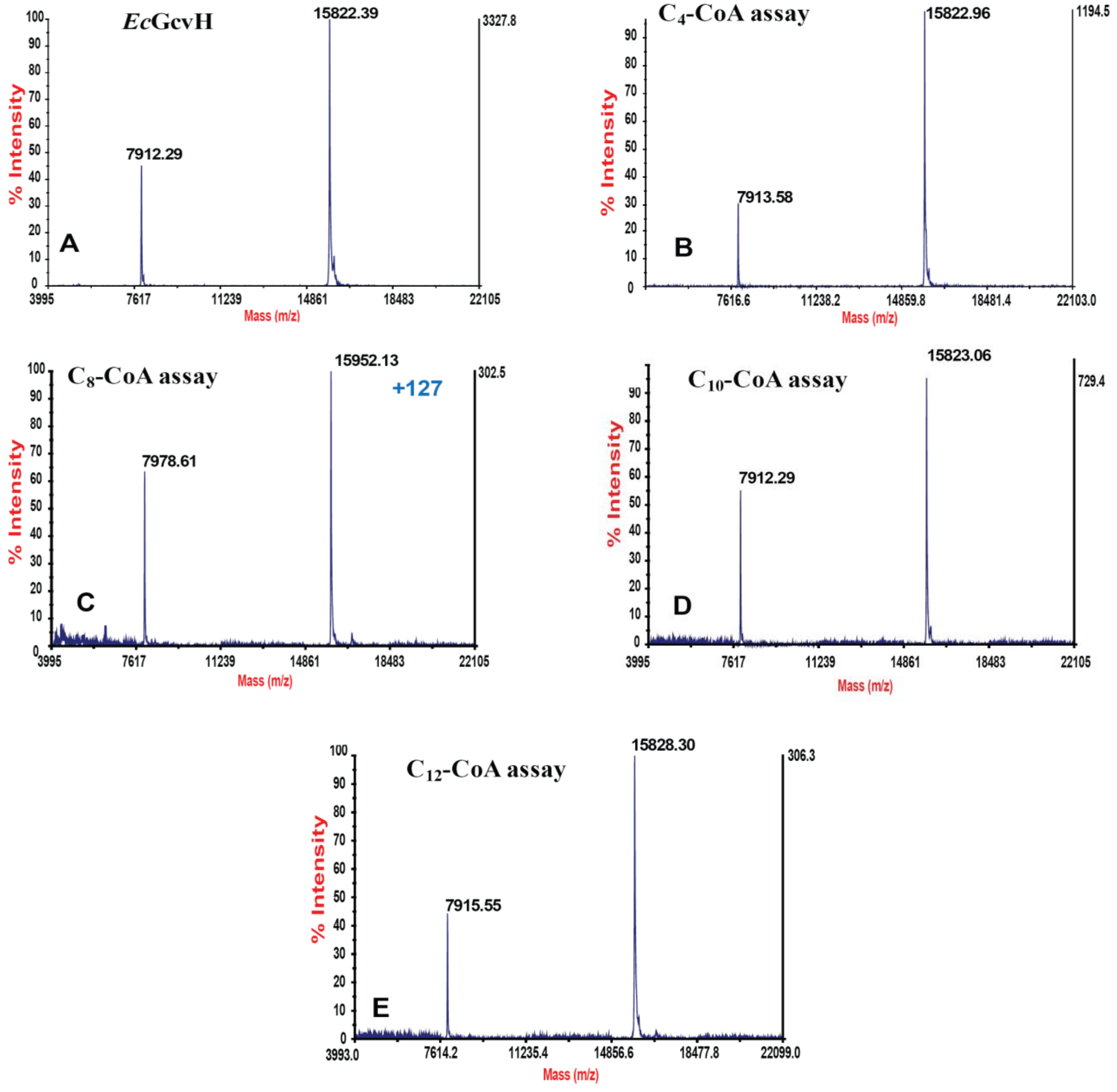


**Fig. S4.** Mass Spectrometry confirms that LipB can transfer octanoyl- chain from $C_8$-CoA to GcvH. MALDI-TOF mass spectra for A) apo- GcvH used as a control (monoisotopic mass 15834Da). GcvH in the assay performed using b) $C_4$-CoA, c) $C_8$-CoA, and d) $C_{10}$-CoA, and d) $C_{12}$-CoA as acyl- chain donors. A 127 kDa increase in mass was observed when $C_8$-CoA was used as a donor.
